# Immunoreactivity against SLC3A2 in high grade gliomas displays positive correlation with glioblastoma patient survival: Potential target for glioma diagnosis and therapy

**DOI:** 10.1101/2023.01.26.525653

**Authors:** Nihal Karakaş, Ozan Topcu, Erdem Tüzün, Özlem Timirci Kahraman, Pulat Akın Sabancı, Yavuz Aras, Berrak Yetimler, Gökçen Ünverengil, Mebrure Bilge Bilgiç, Khalid Shah, Hasan Körkaya

## Abstract

Immunity against cancer cells is at the forefront of fight against highly malignant tumors like glioblastomas (GBM). Autoantibodies and their autoantigen cooperators are one of the promising biomarkers linked to the immune responses and have been reported as having an initiative or prognostic role for certain types of paraneoplastic disorders. Nevertheless, immunoreactivity against antigens expressed in GBM are poorly studied. To date, autoantibodies were identified by pursuing targeted approaches. By contrast, in this study, we collected human GBM tissue and sera samples and by applying proteomics analysis, we determined autoantigen candidates for GBM. Subsequent immunohistochemistry and immunoprecipitation experiments revealed that a solute carrier; SLC3A2 is widely overexpressed in GBMs and therefore, immunoreactivity against SLC3A2 is present in high grade GBM, and SLC3A2 expression is altered in GBMs. No antibody interaction was detected in SLC3A2 expressors of low grade gliomas. Furthermore, autoantibody presence was correlated with prolonged survival of GBM patients. Taken together, for the first time, we reported that the SLC3A2 immunoreaction exists in high grade gliomas with a distinct GBM profile. In conclusion, our findings may open up new avenues for our understanding of glioma prognosis in the context of autoimmunity. This may eventually lead to diagnostic and therapeutic inventions that can be utilized for prevention of the disease progression.

## INTRODUCTION

Glioblastoma (GBM) is an aggressive and malignant brain tumor that is most common in adults [1]. The mean survival for GBM patients is 12-15 months [2]. According to the 2016 report, the World Health Organization (WHO) classified GBM as grade IV among gliomas [3]. Of these, GBMs are then defined as Grad 4 with an IDH wild type phenotype by 2021 [4]. Currently, the standard therapy for GBM is radiotherapy and adjuvant chemotherapy following surgical resection. GBM patients show high resistance to therapy, and the tumor relapses very quickly after surgery. GBM is primarily diagnosed by neuroimaging and tissue biopsy [5]. However, imaging techniques may not accurately distinguish between lesions caused by treatment and those resulting from actual tumor progression [6]. Moreover, tissue biopsies might cause the tumor to become highly invasive [7] and affect neurological functions of patients.

Cancer immunotherapy has been leading the way with remarkable responses in some malignancies such as non-small cell lung cancer (NSCLC) and melanoma [8]. However, it has limited or no activity in majority of cancer patients including those with gliomas. Cancer immunotherapy strategies currently under clinical trials against malignant gliomas involve immune checkpoint inhibitors (ICI), peptide vaccines, dendritic cell vaccines, chimeric antigen receptor T (CAR-T) cell therapies, and oncolytic viruses. Moreover, the discovery of new antigens in personal therapy approaches plays an important role in understanding cancer development and designing patient specific immunotherapies. Tumors, including GBM, usually shed their contents into the circulation and cerebrospinal fluid (CSF) [6]. Tumoral contents containing abnormal molecular changes such as genetic mutations, modifications or overexpressions, which are most common in cancer cells, are recognized as neoantigens by the immune system. Thus, autoantibodies against these neoantigens are produced by plasma cells. Since autoantibodies circulate in the blood, they can be easily detected in patient sera. The determination of the responses by these host immune reactions to the changes undergoing during malignant transformation may potentially enable early diagnosis of GBM. Although anti-neuronal antibody formation triggered by non-neuronal cancers and associated paraneoplastic neurological syndromes have been studied in detail [9], the immunogenicity of the primary central nervous system tumors has been vastly underestimated. Studies conducted in this context have been performed generally against specific proteins or autoantigen candidate peptides that may be related to GBM pathology [7–10]. It is becoming increasingly evident that autoantibodies may contribute to disease mechanisms even when they are not the primary cause of the disease [10], [11]. Therefore, a comprehensive study for autoantibodies produced against neoantigens expressed in GBM tissues is needed.

In this study, our primary goal was to discover autoantibodies specific to GBM. Therefore, we designed our experiments to detect a GBM-specific autoantibody profile by using patient sera instead of primary antibodies while analyzing patient tissues with the WB method. We performed mass spectrometry analyzes for distinct proteins that were specifically present in GBM patient sera. Subsequent mass spectrometry analysis showed that SLC3A2 was overexpressed in GBM tissues compared to epilepsy patients (with no tumor) and non-GBM gliomas. We hypothesized that SLC3A2 overexpression could be perceived as neoantigen by the GBM patients’ immune system. As a result, the presence of SLC3A2 autoantibodies was determined to be specific for patients with in high grade glioma. Moreover, anti-SLC3A2 autoantibody reactivity was found to be positively correlated with the survival of GBM patients.

## MATERIALS AND METHODS

### 1. Sample collection from participants

For this study, 25 patients with glial brain tumors (14 GBM and 11 non-GBM) and 5 epilepsy tissue and 10 healthy volunteers were enrolled by the Department of Neurosurgery, İstanbul Faculty of Medicine, İstanbul University between 2016-2019. All diagnostic criteria of the 2016 World Health Organization Classification of Tumors of the Central Nervous System for glioma and GBM [12]. A standard chemoradiotherapy after surgical resection was applied to all patients [13]. Patients with chronic neurological or systemic diseases other than the brain tumor, those who received immunosuppressant treatment in the last 3 months, and patients younger than 18 were excluded. Tissue and sera samples were stored at −80C° and −20C°, respectively until they were used for experimental purposes. To diagnose the glioma grades, tissue samples of all cases were examined at the Department of Medical Pathology. Medial temporal lobe specimens of epilepsy patients who had undergone epilepsy surgery were used as disease controls. Patient’s survival (days) was calculated as the length time from the diagnosis of GBM to death. The study was approved by the institutional review board and all participants gave their written consents. All procedures performed were in accordance with the ethical standards of the institutional review board and with the 1964 Helsinki declaration and its later amendments.

### 2. Immunohistochemistry

Paraffin embedded brain tissues were prepared, analyzed and diagnosed by Department of Medical Pathology, İstanbul Faculty of Medicine, İstanbul University, Türkiye. Cases were grouped into 3 as control, non-GBM and GBM. After the tissue slides were deparaffinized, they were passed through a 96% −70% alcohol gradient. After blocking with 2% FBS, they were incubated overnight with 1: 3500 SLC3A2 (Sigma / HPA017980). After the washes, incubated with 50 μl of secondary antibody for 30 minutes at room temperature. Tissue samples were washed 3 times for 5 minutes in PBS solution. 50 μl of streptavidin peroxide was applied to the samples and then incubated for 30 minutes at room temperature. Tissue samples were washed 3 times for 5 minutes in PBS and the DAB solution was applied to the samples. Samples were stained with hematoxylin/eosin and passed through 70% −96% alcohol gradient. Samples were put into xylene for 10 minutes, 2 times. The slides are left at room temperature until they dry.

### 3. Protein extraction from human brain tissue

All tissues and sera (n = 20 GB, n = 10 grade III-IV glioma, n = 10 grade I-II glioma, n = 5 epilepsy) were collected by Department of Neurosurgery, İstanbul Faculty of Medicine, İstanbul University, Türkiye. Brain tissues were lysed in RIPA buffer for 30 minutes at 4 C°. The supernatant was collected by centrifugation at 10,000 x g for 10 minutes. The tissue samples were stored at −80 C°.

### 4. Western Blot

40 μg protein samples were run for 1.5 hours at 120 V in 10% polyacrylamide gel and then transferred to PVDF membrane. Membrane was blocked with 5% fatty milk or BSA for 1 hour at room temperature on shaker. Membranes were incubated overnight at +4 C on shaker with human sera (1: 200), ß-actin (1: 5000, Abbkine), α-tubulin (1: 5000, CST), SLC3A2 (0.8: 1000, Sigma). After primary antibody incubation, membranes were washed with 1x TBST for 3×5 min. They were then 1 hour incubated with HRP conjugated secondary antibodies, goat-anti-human 1:3000 (Biorad /1721050); goat-anti-rabbit 1:2000 (CST / 7074); and goat-anti-mouse 1:2000 (Abbkine / A21010) at room temperature on shaker. After the incubation, the membranes were washed 3×5 min with TBST. Next, ECL substrate (Biorad/1705061) added and bands were chemiluminescently detected using Gel Doc (Biorad). Densities of the obtained bands were quantified by a software (Image J).

### 5. Proteomics

#### In gel protein digest

Protein bands from polyacrylamide gels were excised with sharpened micro spatula. The gel was cut into ~1 x 1 mm cubes and placed in 1.5 ml snap-cap microfuge tubes. The gel particles were washed with 100 mM ABC and vortexed briefly, spun down and the liquid was discarded. Next, acetonitrile was added (approximately 3 - 4 times the total volume of the gel pieces) and the gels were incubated for 10 - 15 min until the pieces shrank, became opaque and stuck together. The gel particles were spun down, and after removal of the liquid, they were swelled in 100 mM DTT in 100 mM ABC and incubated for 30 minutes at 56°C to reduce the proteins. This step was repeated with ACN, which was then replaced with 100 mM CAA in 100 mM ABC. After incubation for 20 min at room temperature, the CAA solution was discarded and the gel particles were washed with 100 mM ammonium bicarbonate for 15 minutes. Another step of spinning down and incubation in ACN of the gel particles followed. The rehydrated gel particles were incubated with trypsin overnight in 50 mM ABC at 37 C°. After overnight incubation, all supernatants were collected in fresh tubes and concentrated by speed vacuuming. Tryptic peptides were desalted by Stage Tipping using Empore C18 47mm disks and then dried in Speed-Vacuum. The peptide samples were resuspended in 5% formic acid and 5% acetonitrile for LC-MS/MS analysis.

#### Data acquisition

Peptides were analyzed by C18 nanoflow reversed-phase HPLC (Dionex Ultimate 3000 RSLC nano, Thermo Fisher Scientific) combined with an orbitrap mass spectrometer (Q Exactive Orbitrap, Thermo Fisher Scientific). Samples were separated in an in-house packed 75 μm i.d. × 35 cm C18 column (Reprosil-Gold C18, 3 μm, 200 Å, Dr. Maisch) using 70 minute linear gradients from 5-25%, 25-40%, 40-95% acetonitrile in 0.1% formic acid with 300 nL/min flow in 90 minutes total run time. The scan sequence began with an MS1 spectrum (Orbitrap analysis; resolution 70,000; mass range 400–1,500 m/z; automatic gain control (AGC) target 3e6; maximum injection time 60 ms). Up to 15 of the most intense ions per cycle were fragmented and analyzed in the orbitrap with Data Dependent Acquisition (DDA). MS2 analysis consisted of collision-induced dissociation (higher-energy collisional dissociation (HCD)) (resolution 17,500; AGC 1e6; normalized collision energy (NCE) 26; maximum injection time 100 ms). The isolation window for MS/MS was 2.0 m/z.

#### Data processing

Peptide identification and quantification from raw MS data files was done using the MaxQuant software (v1.6.0.1) [14]. Peptide spectral matches were searched against a Swissprot database containing 21,039 entries for *Homo sapiens* retrieved from Uniprot in 2016. Trypsin was selected as hydrolytic enzyme with a maximum number of allowed missed cleavages of 2. Methionine oxidation and protein N-terminal acetylation were set as dynamic modifications, while cysteine carbamidomethylation was set as fixed modification. The False Discovery Rate (FDR) for peptide and protein identifications was set to 0.01. Only peptide identifications with a sequence length between 7 and 25 were allowed. Matching between runs was allowed for fractions of the same sample type with default settings. All other settings were default.

#### Bioinformatic Analysis

The proteome data was searched against the reviewed human protein database from Uniprot. Similar proteins were grouped and quantitative value was given for the one with the highest score. *p* values and experimental ratio of proteins were calculated and filtered according to these two parameters for further experiments. Gene ontology analysis (GO) provides functionally annotation of DEGs within a biological context with respect of biological processes, cellular components and molecular functions [15]. Kyoto Encyclopedia of Genes and Genomes (KEGG; http://www.kegg.jp/) pathway database was used for analyzing gene functions and to link this data with molecular pathways [16]. GO terms and KEGG pathway enrichment analyses for DEGs were performed by using the database for annotation, visualization and integrated discovery (DAVID) online database (DAVID Bioinformatics Resources 6.8). Search Tool for the Retrieval of Interacting Genes/Proteins (STRING; http://www.string-db.org/) is a database that identifies known and predicted interactions between proteins [17]. Selected pathways were filtered into the DEG protein–protein interaction (PPI) network complex containing nodes and edges with parameters including a minimum required interaction score > 0.4 (medium confidence). Only query proteins were displayed. Hub genes, highly interconnected with nodes in a module, have been considered functionally significant. A hub is a node that has a higher degree than other nodes in the graph. In our study, hub genes were defined by module connectivity. Furthermore, we uploaded all genes in the hub module to the STRING database to construct PPI to screen hub nodes in PPI network. We defined genes with the node connectivity > 2 (total edges/total nodes) as the hub nodes in PPI network. Additionally, the genes with p value less than 0.05 were identified as real hub genes.

### 7. Immunoprecipitation of SLC3A2 from tissues

In our study, SLC3A2 was precipitated using anti-SLC3A2 antibodies (Sigma/HPA017980) from tissue lysates for WB analysis. After imaging the membranes incubated with patient sera, the membrane was stripped and incubated again with anti-SLC3A2 primary antibody to ensure loading control. Briefly, the next immunoprecipitation protocol was applied to carry out the study in this direction. 20 μl A/G sepharose beads (GE Healtcare/ 17-0618-01) were added onto 500 μg lysate and incubated at +4 C for 1 hour at 25 rpm. Samples were centrifuged at 3000 rpm for 2 minutes and the supernatant was taken into clear tube. 1 μg of anti-SLC3A2 were treated on the supernatant for 1 hour at +4 C and 25 rpm. 30 μl A / G sepharose beads were added to the mixture and incubated at +4 C and 25 rpm overnight. After being centrifuged at 3000 rpm for 2 minutes, the pellet was washed 2 times with 1 ml 1x PBS (Gibco / 10010023). After it was centrifuged at 3000 rpm for 2 minutes and the supernatant was discarded, the pellet was treated with 10 μl 4x Laemli buffer (Biorad /1610747) and boiled at 100 ° C for 10 minutes.

### 8. Immunofluorescence Assay

LN229 (ATCC / CRL-2611), T98G (ATCC / CRL-1690) and U87 GBM (ATCC / HTB-14) cells were cultured with high glucose DMEM (Gibco) and %10 FBS (Gibco). LN229 GBM cells were seeded into 6-well plates at 3-5 x 10^5^ cells. After fixation with 4% PFA, cells were washed 2 times with PBS. 500 μl of blocking solution (Thermo / 37520) was added to each petri dish and incubated at room temperature for 1.5 hours. Then the wells were washed 2 times with PBS and 200 μl of SLC3A2 (1: 500) (Sigma / HPA017980) primary antibody solution (in 5%BSA) was added to each petri dish and incubated overnight on a shaker. After washing with PBS 2 times, 200 μl of 1: 500 diluted secondary antibody solution (Alexa Fluor Plus 488, Thermo) (in 5%BSA) was added and incubated for 1.5 hours at room temperature on a shaker. Secondary antibody was discarded and the plates were washed with PBST 4 times for 4 minutes on a shaker. To stain nuclei, 500 μl of DAPI was added in each petri dish and kept on a shaker for 3 minutes. After washing with PBST for 5 minutes, next washed once with PBS.

### 9. Statistical analysis

Data analysis was performed using the Perseus platform (v1.6.2.2) [18]. Peptides only identified by sites, by reverse sequence and as potential contaminants were removed from the final data. Statistical analysis for the determination of differentially regulated proteins was conducted by applying two-sided two-sample Student’s *t*-test between the GBM vs Control and GBM vs non-GBM samples. Proteins with a p-value <0.05 were considered as significant. Statistical tests were carried out using an unpaired t-test for WGCNA. All analyzes were carried out using the Graphpad program. One Way and Two Way ANOVA was used for immunoblot and immunoprecipation analysis, respectively. P=**** <0.001 and p=***>0.001 and P=* <0.05 were considered as significant. Correlation analysis was done with Spearman’s correlation test.

### 10. Ethical Issues

This study was approved by İstanbul Medipol University, Non-Interventional Clinical Research Ethics Committee Presidency (no: 10840098-604.01.01-E.6985).

## RESULTS

### 1. SLC3A2 is differentially expressed in GBM tumor

To examine whether there is a specific immunoreactivity between GBM patient sera and GBM tissue, samples were obtained from glioma cases (grad I to grad IV) as well as epilepsy patients (without tumor diagnosis) as control cases. Sera of patients were collected prior to neurosurgery. In the WB experiments, case sera were used instead of primary antibodies. GBM sera showed a distinctive band pattern with GBM tissue more visibly around 50kDa, then the identical protein-sera interaction was determined and applied for further proteomics analyses (Fig. 1A).

**Figure 1.**
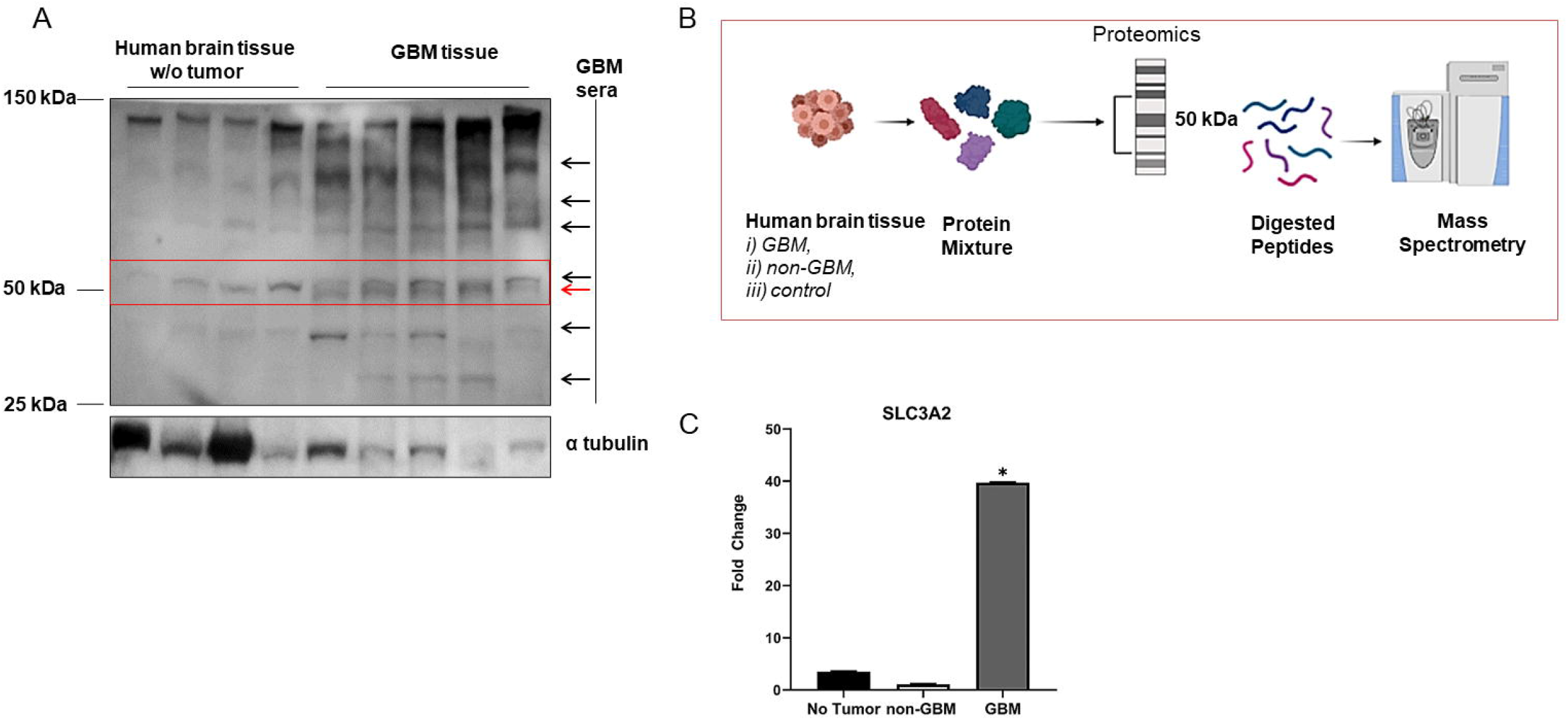
GBM sera selectively interacts with GBM tissue and immunoreactions suggest SLC3A2 as a potential auto-antigen. A) Lysates from GBM and epilepsy (no tumor) tissues were run on SDS-PAGE and western blotted using GBM patient sera instead of primary antibodies (no significant change was recorded when using purified IgGs from sera) and α tubulin was used as the loading control. B) Tissue lysates were analysed by mass spectrometry (p=*<0.05) regarding the autoantibody-autoantigen interactions. C) SLC3A2 expression is 11 times folded as compared to control tissue lysates in GBM (p=*<0.05).

Mass spectrometry analysis was performed with lysates obtained from 5 control, 5 non-GBM glioma and 5 GBM patient tissues (Fig. 1B) yielding identification of 3014 potential protein candidates. Proteomics analyses were performed using the “Thermo Orbitrap Exactive-Bottom up” strategy. Max Quant and Perseus programs were used for the first analysis (p<0.05, fold change≥2). DAVID (GO and pathway analysis) and String (protein-protein interaction) programs were used for bioinformatics analysis. GO analysis revealed that differential expression of proteins were involved in 1 biological processes, 1 cellular component and 1 molecular function (Table 2). KEGG pathway analysis indicated that upregulated proteins were mainly enriched in spliceosome, alcoholism, systemic lupus erythematosus, rna transport, ribosome, amoebiasis, protein digestion and absorption (data not shown). Data for control and non-GBM gliomas were compared with GBM patients by bioinformatic analyses. Accordingly, 10 proteins were found to be statistically significant and no interaction was found depending on the String analysis. Of these, SLC3A2 expression was 39.8 and 11-fold higher compared to non-GBM gliomas and epilepsy control group, respectively (Table 3, and Fig. 1C). To determine possible antigen candidates, we focused on the proteins around 50 kDa since it was distinctly different from non-Glioma controls (Fig. 1A). Accordingly, 3 differentially expressed proteins were considered in that respect and no interaction was detected with these antigen candidates and patient sera (data not shown).

**Table 1.** Average age and gender distribution of study groups.

| Tumor | Age (Mean) | Gender (M/F) |
| --- | --- | --- |
| Glioblastoma Multiforme (GBM), n=14 | 61.07 ± 12.88 | 10/4 |
| Grade III-IV Glioma, n=6 | 51.2 ± 12.09 | 3/3 |
| Grade I-II Glioma, n=5 | 39 ± 5.78 | 4/1 |
| Epilepsy, n=5 | 21.6 ± 10.59 | 3/2 |
| Healthy controls, n=10 | 27 ± 4.97 | 4/8 |

SD, standard deviation; M, male; F, female

**Table 2.**
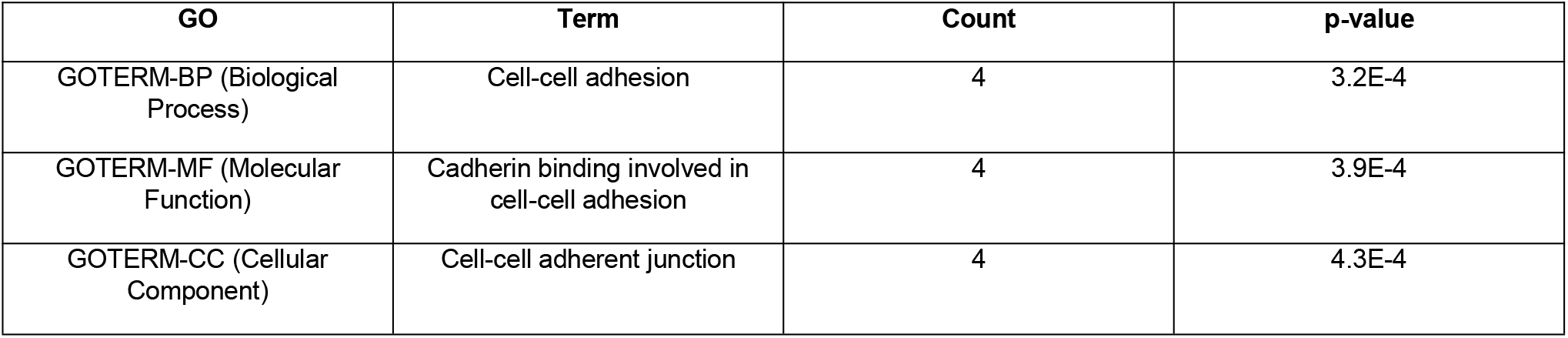
GO Analysis of Differentially Expressed Proteins

| GO | Term | Count | p-value |
| --- | --- | --- | --- |
| GOTERM-BP (Biological Process) | Cell-cell adhesion | 4 | 3.2E-4 |
| GOTERM-MF (Molecular Function) | Cadherin binding involved in cell-cell adhesion | 4 | 3.9E-4 |
| GOTERM-CC (Cellular Component) | Cell-cell adherent junction | 4 | 4.3E-4 |

**Table 3.** SLC3A2 expression metrics in patients with GBM versus non-GBM based on the identification by LC–MS/MS

| Protein IDs | Protein name | Gene name | Fold Change | MW [kDa] | p-value |
| --- | --- | --- | --- | --- | --- |
| P08195 | 4F2 cell-surface antigen heavy chain | SLC3A2 | 39,802 | 67,993 | 9,56267E-08 |

### 2. High grade gliomas develop anti-SLC3A2 autoantibodies

Immunoprecipitation studies were performed to detect the presence of autoantibodies in GBM sera against SLC3A2, which is found to be overexpressed in glioblastoma tissues. To further confirm, control (n=5), non-GBM (n=7) and GBM (n=14) tissue lysates were precipitated with commercial SLC3A2 antibody and analyzed using GBM sera (n=10) as a primary antibody in WB analyses (Fig 2A). Expectedly, patient sera interacted strongly with high grade glioma tissue lysates while it interacted weakly with grade I-II glioma tissue (Fig 2B). No interaction was observed with grade I-II glioma sera (n=5) (data not shown). Immunoprecipitation studies were also performed using non-GBM patient sera to determine whether autoantibodies in GBM sera were present in other gliomas. It was observed that high grade non-GBM sera (grades III and IV, n=5) showed weak interaction with GBM tissue lysates compared to GBM sera. Low grade sera (grades I and II, n=5) showed no detectable interaction with tissue lysates. Interestingly some of the GBM patient sera showed specific reaction with GBM tissues only (Fig 2B). Therefore, the significance of both GBM and non-GBM patient sera interaction with SLC3A2 were analyzed. It was found that GBM sera showed greater (43 times more) reactivity with SLC3A2 in GBM tissue as compared to non-tumor group (Fig 2C). It was determined that non-GBM high grade glioma (grade III-IV) sera showed 34.5 times higher reactivity with SLC3A2 in GBM tissue as compared to non-tumor group. Moreover, GBM sera interaction with SLC3A2 in GBM tissue was significantly higher than non-GBM high grade glioma sera (Fig 2C). The results demonstrated that high grade gliomas develop immunoreactivity against SLC3A2 and thus GBM sera showed the strongest reactivity to the SLC3A2 in GBM tissue compared to grade III-IV sera.

**Figure 2.**
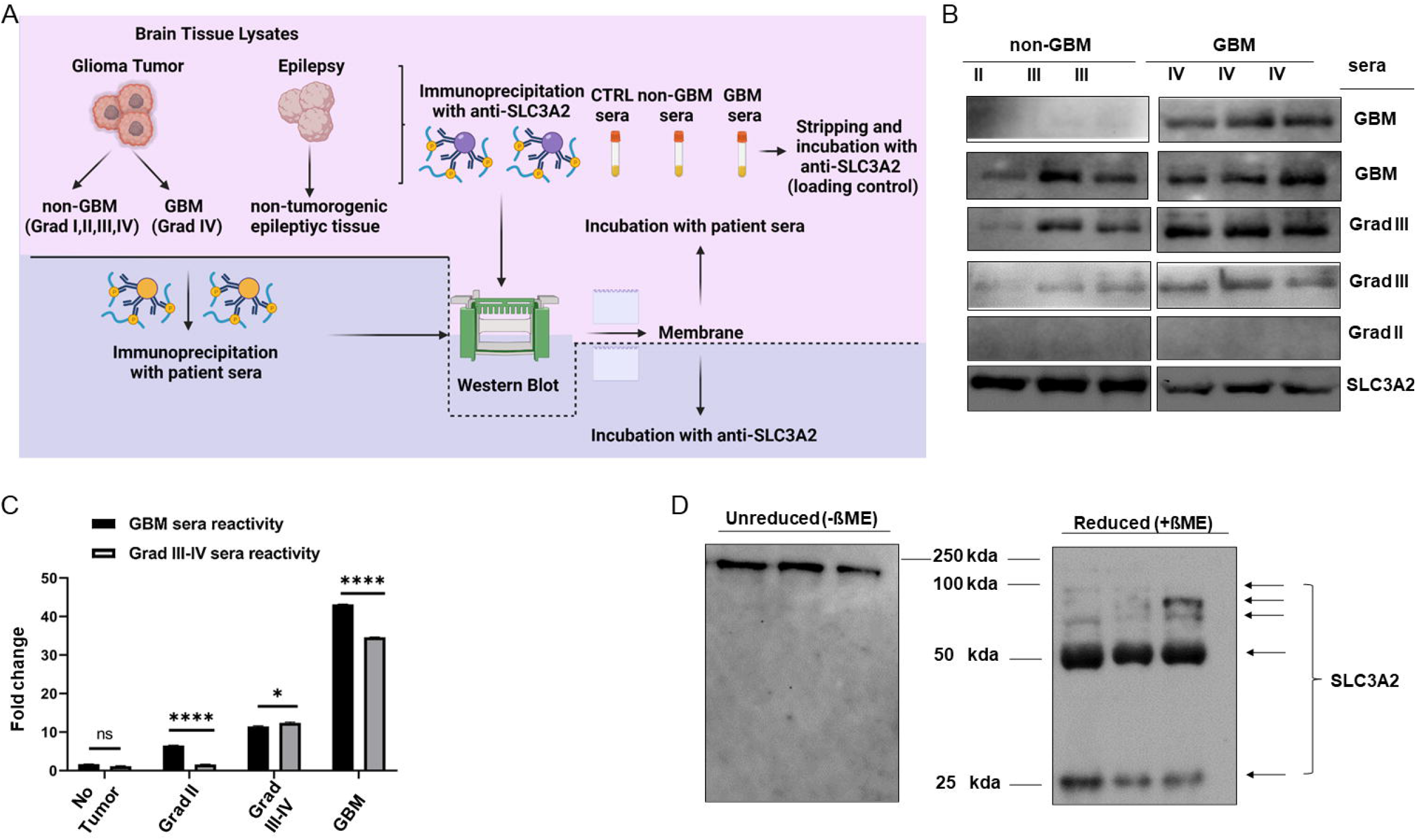
Patient sera from high grade gliomas showed immunoreactions against SLC3A2. A) Schematic representation of immunoprecipitations. GBM (n=14); non-GBM glioma (n=8); and epilepsy tissues (with no tumor as of control), (n=5) were used as protein sources for SLC3A2 precipitatation with commercial SLC3A2 antibody. In a set of experiments patient sera was also used for SLC3A2 extraction from the tissues. Western blotting was applied to the mebranes using patient sera (n=10 GBM sera; n=5 high grade gliomas’ sera; n=5; low grade sera; n=10 healthy control sera). Then the stripped membranes were incubated with SLC3A2 antibody to ensure accurate loading of SLC3A2 precipitates for each group. B) Immunoreactivity against SLC3A2 in GBM and non-GBM tissues in the presence of sera from varying glioma grades. No interaction was observed with the control sera applied group (data not shown). C) ImageJ analysis of band densities indicated that sera of high grade gliomas showed increased reaction against SLC3A2, and GBM sera interaction with SLC3A2 in GBM tissue was significantly higher than non-GBM high grade glioma sera. (Two Way ANOVA, p=*<0.05;****<0.0001). D) Reduced (in the presence of ß-mercaptoethanol) and unreduced GBM tissue lysates were used for SLC3A2 precipitation by GBM sera. Bands depict 3D binding of the antibody and the antigen couple in unreduced samples while reduced samples showed antibody interaction with different forms of the SLC3A2 in GBM tissue.

To further characterize the antibody development against SLC3A2 in GBMs, 3D binding nature of the autoantibody was considered since immunoprecipitated SLC3A2 in wetern blot analyzes was linearized. The data from those studies indicated that anti-SLC3A2 autoantibodies actually bind linear form of SLC3A2. To clarify, whether anti-SLC3A2 autoantibodies in GBM sera can recognize the 3-dimensional structure of SLC3A2 (considering the antibody development in vivo), they were analysed with or without beta-mercapto ethanol (ßME). Our findings revealed that autoantibodies in GBM sera can bind to the 3D structure of SLC3A2 in GBM tissue as well as to the linearized form. Molecular weight of antibody-antigen binding was found to be correlated with the total mass of the couple. (Fig 2D).

### 3. SLC3A2 immunoreactivity displays subspesific profile in GBM patients

Immunoprecipitation studies revealed that GBM sera interacted strongly with SLC3A2 in GBM tissues while it was a weak interaction in non-GBM gliomas and control tissues. In addition, certain type of GBM sera distinctively interacted with GBM tissue, by contrast not with other high grade gliomas (Fig 3A-B). The difference in this interaction pattern was considered as variations of the antigen;SLC3A2 in GBM tissues since the SLC3A2 in non-GBM gliomas and control sera was present on the same membrane incubated with GBM sera (incubation with SLC3A2 antibody depicted accurate SLC3A2 precipitations for each glioma sample on stripped membranes). Accordingly, common SLC3A2 isomers and modifications were searched using the UniProt database and thus ten SLC3A2 isoforms were found to be available. Of these, 3 SLC3A2 isoforms were in the range of molecular weights we determined in our study (H0YFS2, 25 kda; F5GZS6; 64 kda, J3KPF3, 68 kda). Among these 3 candidates, 2 isoforms (in the range of 30-70 kDa) with database entry names F5GZS6 and J3KPF3 were correlated with the interactions we spesifically detected (around 50kDa) (see Fig 1A). Interestingly, various amino acid modifications of SLC3A2 were defined in the UniProt database. When these modifications were examined, SLC3A2 phosphorylation was determined at multiple residues of SLC3A2, more frequently at serine residues. To study possible modifications of SLC3A2 in GBM tissues, we then used phospho-serine antibody in WB analyses following previously explained SLC3A2 precipitation methods. Together our studies revealed that SLC3A2 in GBM tissues was found to be approximately 2.5 times more phosphorylated compared to non-GBM and control tissues, while there was no significant difference between non-GBM gliomas and the control tissues (Fig 3C-D).

**Figure 3.**
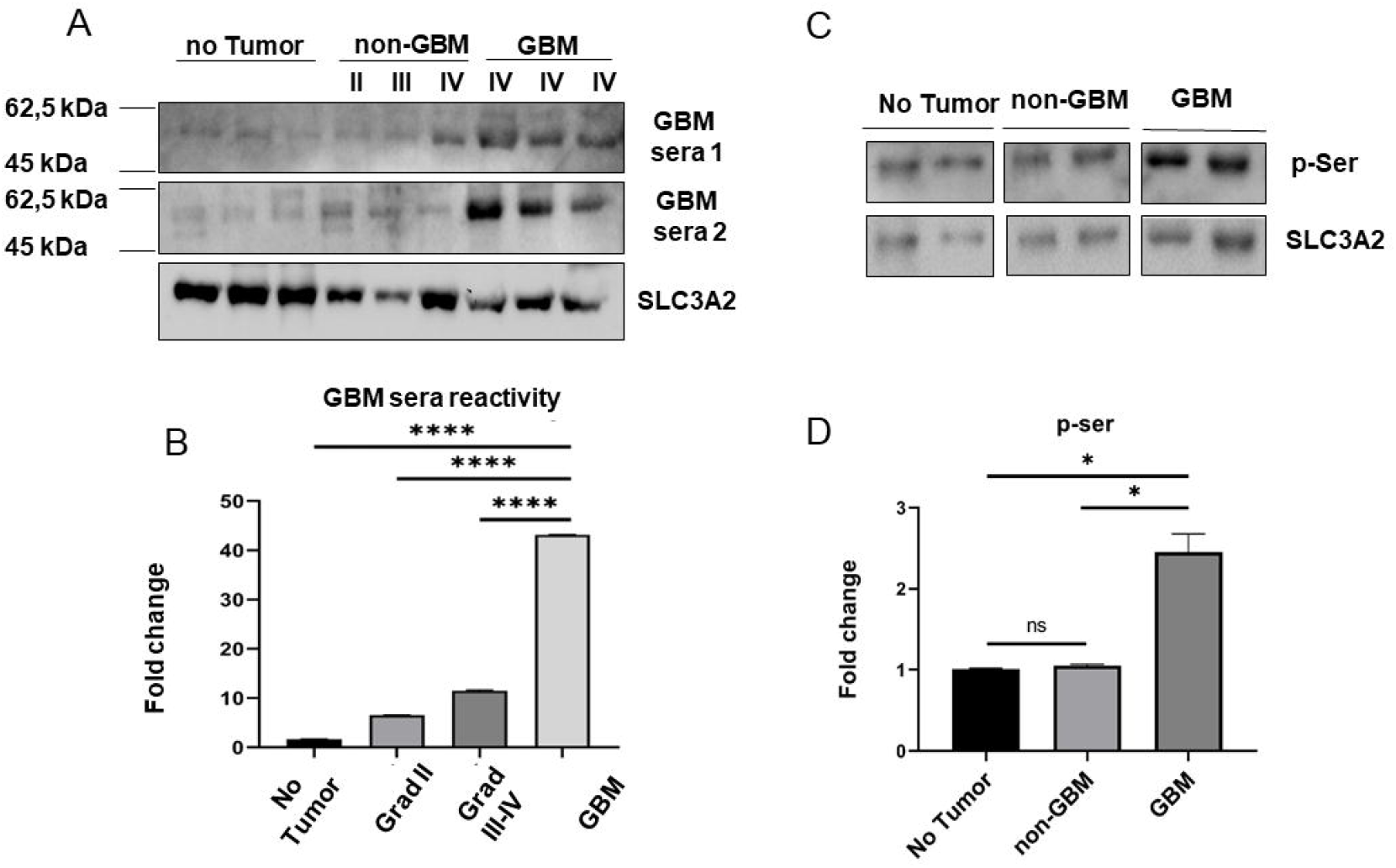
Glioblastoma patients display GBM specific SLC3A2 immunoreactivity. A) Western blot of immunoprecipitates using commercial SLC3A2 antibodies, and immunoblots with GBM sera as of primary antibody. Commercial SLC3A2 primary antibody was used as a loading control for SLC3A2 precipitation upon membrane strips B) GBM sera reactivity was accounted by band intensities. GBM sera (n=10) showed approximately 43, 5 and 9 times more strong reactivity compared to the control sera; grad III-IV glioma (n=5); and grad I-II glioma (n=5), respectively (One Way ANOVA, p=****<0.0001).) C) Immunoprecipitation with SLC3A2 antibody and immunoblotting with phospho-Serine (p-Ser) antibody demonstrates the phosphorylation status in GBM (n=9); non-GBM (n=3) and control (n=3) tissues. SLC3A2 levels in precipitates were used as loading control on stripped membranes. D) SLC3A2 in GBMs was counted approximately 2.5 times more phosphorylated compared to non-GBM gliomas as well as the control group (One Way ANOVA, p=*<0.05).). In all control groups no tumor bearing epilepsy patient tissue was used.

### 4. SLC3A2 in GBM cell line apperantly interacts with GBM patient sera

To investigate how GBM cells behave *in vitro* when exposed to autoantibody against SLC3A2 involving patient sera, first, we analyzed SLC3A2 expression levels in GBM cell lines. We found that LN229 and T98G GBM cell lines express significantly increased levels of SLC3A2 compared to U87 GBM cell line (Fig 4A). Next, we established 5 experimental groups to image the SLC3A2-specific binding affinity of anti-SLC3A2 autoantibodies in GBM patient sera. In these experimental groups, the sera of GBM (n=5), grade III-IV non-GBM glioma (n=5), garde I-II (n=5) non-GBM glioma and control (n=5) cases were pooled separately for each group. LN229 cell lines were incubated with the sera of each group for 24 hours. Negative control group without sera incubation was also included. After 24 hours of sera incubation, cells were examined for SLC3A2 binding by using immunofluorescence staining. SLC3A2 staining was significantly decreased in the groups incubated with GBM sera (Fig 4B). These results were correlated with our previous findings with immunoprecipitation experiments. Therefore, we showed that sera incubation induced a significant decrease in SLC3A2 signal intensity in GBMs, suggesting that autoantibody binds to SLC3A2 (Fig 4C).

**Figure 4.**
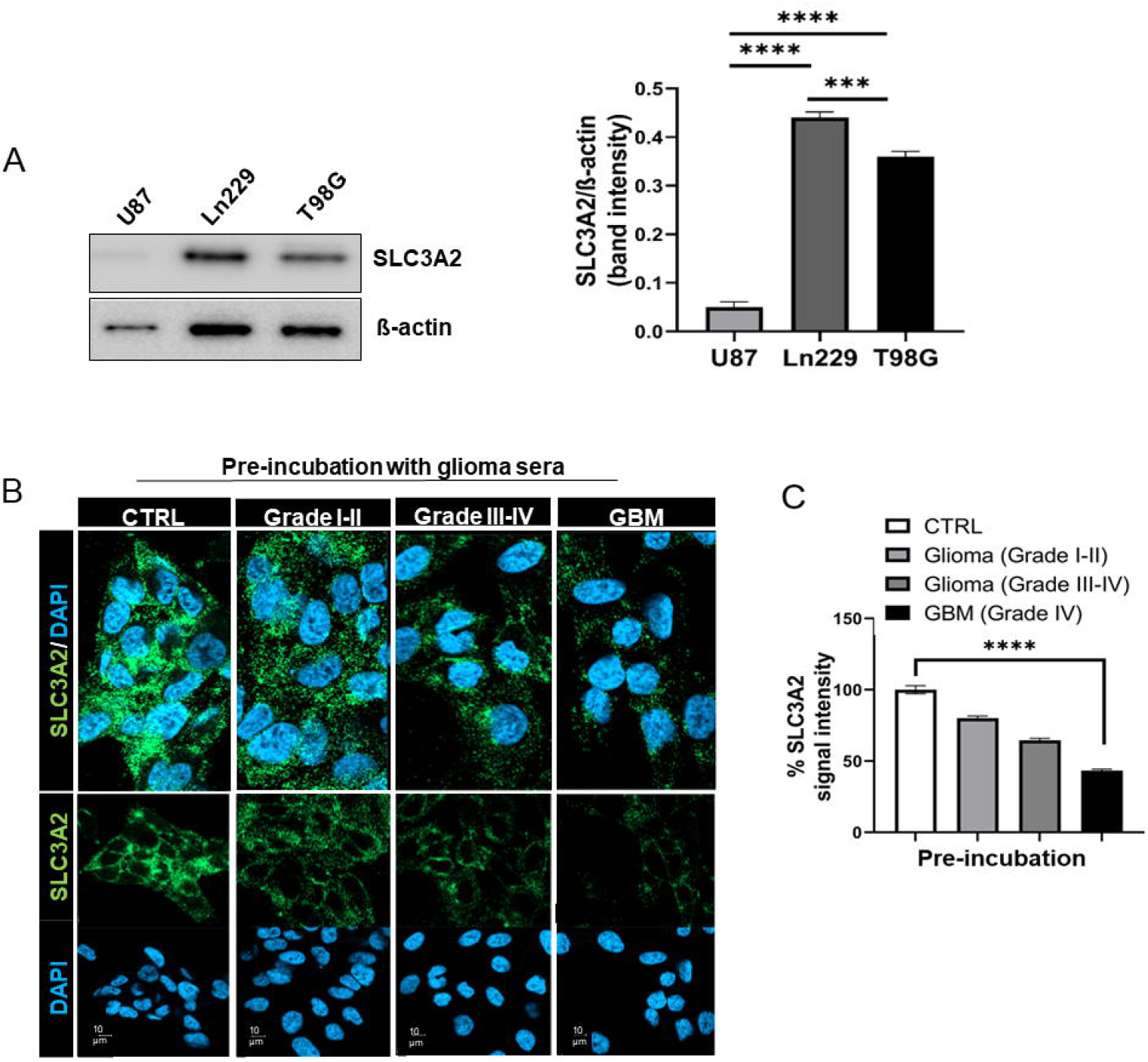
Autoantibodies display SLC3A2-spesific binding profile. A) SLC3A2 expression levels of GBM cell lines (U89, Ln229, and T98G) and Image J analysis of relative band densities. (One Way ANOVA, p=***>0.0001, p=****<0.0001).)B) Ln229 cells were pre-incubated with GBM; grade I-II glioma; grade III-IV glioma; and control sera for 24 hours (n=3 for each group) and SLC3A2 binding to commercial SLC3A2 antibody was visualized for each group (green color indicates SLC3A2 binding in either control or sera preincubatied cells) C) Plot indicates image processing of SLC3A2 for signal intensity. SLC3A2 binding affinity was significantly decreased in GBM sera preincubated cells compared to the control sera (cell were preincubated with healthy sera in control groups). (One Way ANOVA, p=****<0.001).

### 5. SLC3A2 autoantibodies predict prolonged survival in GBM patients with

To determine the link between autoantibody presence and GBM patient survival, we determined protein densities of auoantibody-autoantigen immunoreactions. GBM patients showed varying antibody dilutions and the patients’ survival were detected longer in sera with increased autoantibody. These outcomes indicate the positive correlation between GBM patient survival and SLC3A2 autoantibody presence (Fig 5).

**Figure 5.**
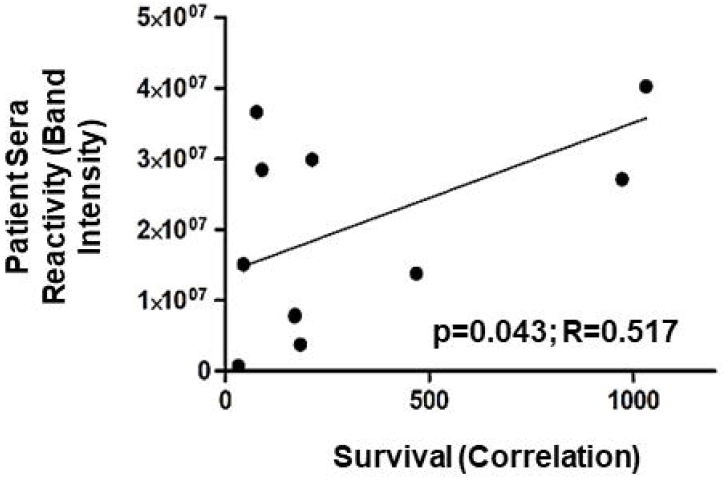
Patient sera reactivity shows positive correlation with GBM patient survival (p=*<0.05). Band intensities of antibody-antigen interactions obtained from immunoprecipitations were analyzed by ImageJ and each dot indicate individual autoantibody densities. The graph shows that the concentration of autoantibody against SLC3A2 is positively correlated with the prolonged GBM patient survival.

## DISCUSSION

Glioblastoma (GBM) is a malignant brain cancer with a median survival of 12-15 months after diagnosis. Due to high mortality rate among GBM patients, there is an unmet need of developing early diagnostics and effective treatment options. Recent advances suggested that autoantibodies may be utilized for early diagnostics and therapeutic responses. However, current studies widely involve targeted approaches focusing on already defined proteins in the pathology. Thus far, GBM-specific autoantibodies enabling the discovery of novel autoantigens have not yet been reported. In this study, glioma patient sera, stratified based on the latest WHO classification during sample collection, were analyzed for possible autoantibodies. Immunoprecipitates obtained by incubation of sera with glioma or epileptic brain tissue were subjected to proteomics analysis. GBM-specific proteins with highest scores were further investigated as potential target antigens and a solute carrier protein SLC3A2 exhibiting a 10-fold expression was selected as the candidate GBM-specific autoantigen. Furthermore, a positive correlation was found between anti-SLC3A2 levels and survival time of GBM patients suggesting potential involvement of SLC3A2 in GBM development and indicating this protein as a potential target for anti-GBM therapeutic research.

SLC3A2 (4F2; CD98) is responsible for polyamine (PA) transport in mammalian cells. SLC3A2 performs its transport task by forming a complex with SLC7A5 (LAT1). LAT1 is overexpressed in tumor cells and thus in pre-clinical treatment for potential anticancer drug target [19]–[21]. SLC3A2 is responsible for the stabilization of LAT1 in this heterodimeric complex and its correct localization on the cell membrane [20]. SLC3A2 / LAT1 expression is very closely related to the proliferation of gliomas and malignant phenotypes [22]. In the last decade, SLC3A2 has been shown to be involved in chemoresistance of cancer cells, and a biomarker for type II renal cancer [23]. To the best of our knowledge, our study is the first to establish a link between anti-SLC3A2 antibodies and glial tumors.

We show that increased expression of SLC3A2 is not only found in GBM specimens but also in glial cell lines LN229 and T98G. While SLC3A2 expression is mostly confined to vascular structures in the non-GBM tissue, GBM tissue exhibits a more widespread expression pattern and a significantly higher expression level. More importantly, several immunoprecipitation studies with patient sera and tumor tissue lysates showed that SLC3A2 protein and autoantibody interactions are specific to high grade gliomas. The binding affinity was significantly pronounced in grade III-IV gliomas while no interaction was observed with sera of grade II glioma patients. Also, sera of some GBM and Grade III/IV glioma patients showed immunoreaction with tissue lysates of GBM but not high-grade glioma specimens. These results were most likely not due to reduced expression of SLC3A2 in low grade gliomas since immunoprecipitation studies done with a commercial SLC3A2 antibody showed that SLC3A2 was expressed in both low- and high-grade glioma tissues and equally precipitated. Finally, incubation of GBM (but not non-GBM) patient sera with SLC3A2-expressing glioma cell lines covered most of the SLC3A2 epitopes, almost eliminating binding sites of the commercial anti-SLC3A2 antibody.

Thus, the distinct interaction profile of anti-SLC3A2 with GMB tissue might be due to the presence of specific SLC3A2 isoforms, post-translational modifications or three-dimensional conformations of SLC3A2 expressed by the GBM tissue. In line with our findings, alterations in post-translational and conformations have been shown to affect the binding of other autoantibodies such as the antibodies of NMDA receptor encephalitis patients [24, 26].

In this context, the Uniprot data showed that SLC3A2 was highly phosphorylated at serine residues in GBM. Accordingly, we could detect significantly increased levels of phosphorylated SLC3A2 in GBM tissues, suggesting that this feature could be one of the potential causes of distinct anti-SLC3A2-GBM interaction. Further investigation of the underlying cause of the distinguishable pattern of SLC3A2 expression in GBM may provide a new perspective for understanding the pathophysiology of the disease. These studies may then lead to the definition of a subset of GBM or high-grade gliomas. On the other hand, if autoantibodies against the SLC3A2 can recognize this unique form of SLC3A2 expressed by GBM, this might then lead to a broad range of therapeutic/diagnostic interventions.

Importantly, our findings considering 3D binding nature of the antibody-antigen highlighted that autoantibody against SLC3A2 developed in GBM patients was specific for SLC3A2 only, not for the complex structures that SLC3A2 can form intracellularly (such as LAT1 association). However, the observation of several bands in the ßME treated samples indicated that anti-SLC3A2 autoantibodies in GBM sera may recognize several SLC3A2 isoforms available on uniprot database.

Another intriguing finding in our study was the positive correlation between anti-SLC3A2 immunoreactivity and survival of GBM patients. These results indicates that GBM patients with higher SLC3A2 immunoreactivity were more likely to display longer survival time. SLC3A2 is over-expressed by GBM and is a part of an aminoacid channel complex that is essential for the survival of GBM cells [27]. Therefore, we reasoned that SLC3A2 antibodies interfere with the functions of SLC3A2 on the cell membrane and limit the nourishment of GBM cells, thereby restricting the growth of the GBM tumors. This assumption needs to be further corroborated by mechanistic studies in future studies.

Taken together, our results reveal two fundamental findings: 1) High grade glioma patients develop SLC3A2 autoantibodies, immunoreactivity of which correlate with survival time. 2) SLC3A2 appears to display a unique expression profile in GBMs. This feature needs to be further studied in future studies for its putative diagnostic and prognostic biomarker potential in GBM patients. Thus, these results provide a compelling evidence for the autoimmunity-glioma interplay in the physiopathology of GBM. Whether there is a subgroup of SLC3A2-antibody positive GBM patients with distinctive prognostic features also needs to be further scrutinized.

Collectively, overall findings open up new discussions on SLC3A2 oriented glioma diagnosis and therapy options through a 4P medicine perspective (Fig 6).

**Figure 6.**
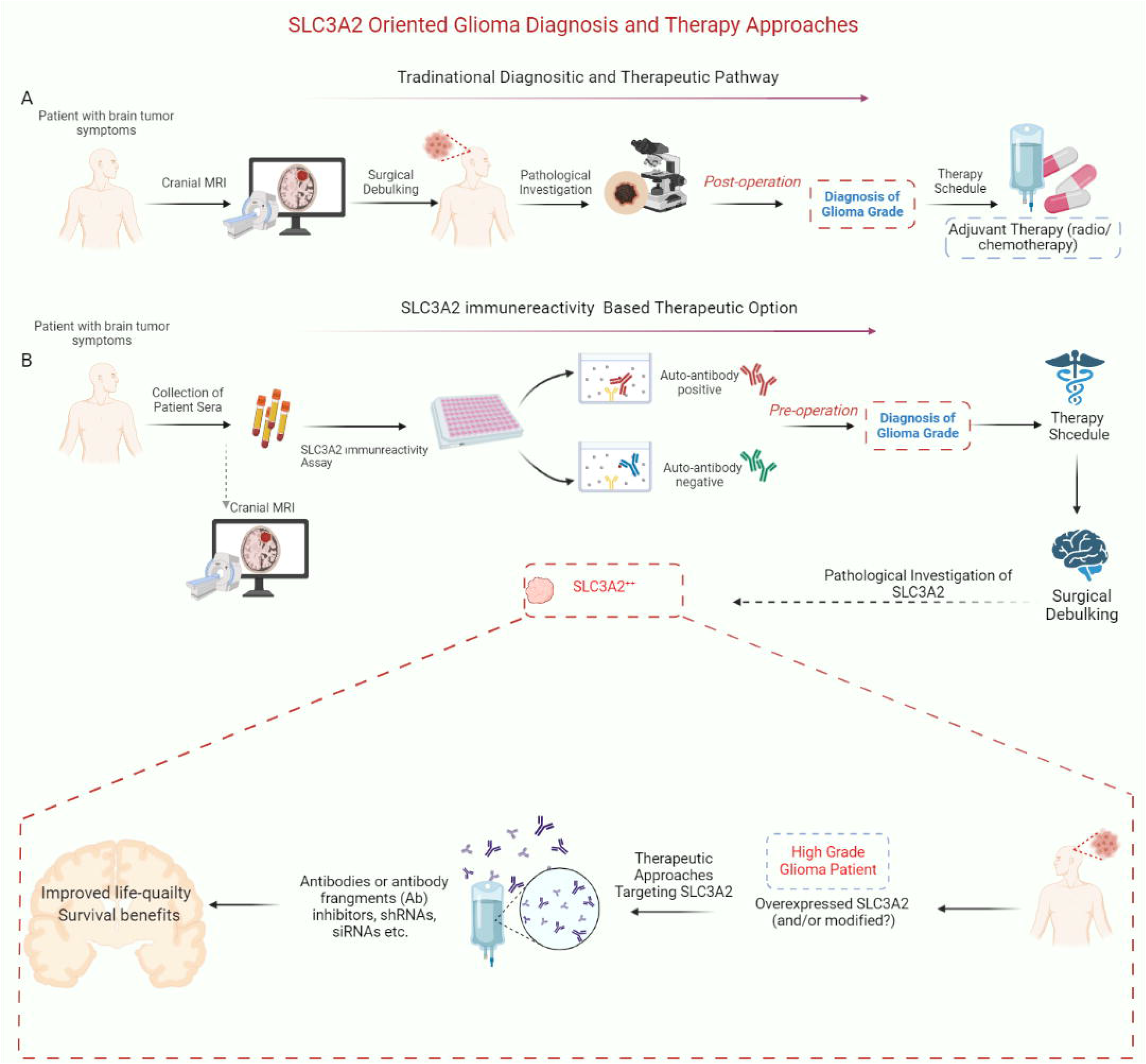
Schematic representation of SLC3A2 immunoreaction orchestrated glioma diagnosis and therapy options. A) Current diagnosis and therapy schedule for gliomas. B) Individualized route for glioma diagnosis and therapy based on SLC3A2 overexpression

## ACKNOWLEDGEMENTS

This work was supported by TÜBÌTAK with project number 216S358 (N.K.). We would like to thank to Aydanur Şentürk for her help with proteomics analysis. We would like to also thank to COST action CA17104 for supporting O.T. with short-term scientific mission (STSM).

## AUTHOR CONTRIBUTIONS

N.K.; conceptualization, funding, design of the study, data assembly, investigation, methodology, supervising, writing-reviewing-editing and final approval of the manuscript, O.T.; design of the study, data collection and assembly, investigation, methodology, formal analysis, writing of the manuscript, final approval of the manuscript, E.T.; conceptualization, design of the study, investigation, methodology, writing-reviewing-editing and final approval of the manuscript. Ö.T.K.; data collection and assembly, investigation, formal analysis, writing and final approval of the manuscript, P.A.S., Y.A.; sample collection, investigation, formal analysis, reviewing and final approval of the manuscript, B.Y.; sample collection, investigation, reviewing and final approval of the manuscript, G.Ü., M.B.B.; sample and data collection, formal analysis, reviewing and final approval of the manuscript, K.S., H.K.; Reviewing/editing and final approval of the manuscript, All authors have read and agreed to the published version of the manuscript.

## Supplementary Figures

**Supplementary Figure 1.**
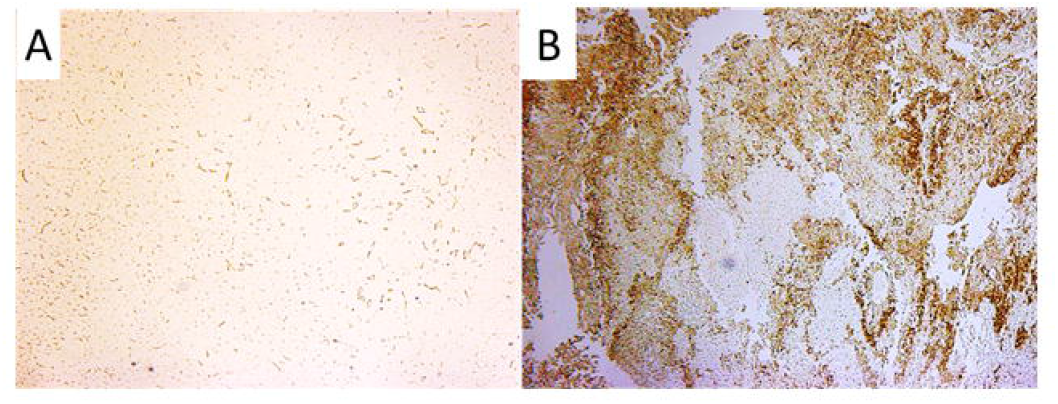
Immunohistochemical expression pattern of SLC3A2 in Grade II (non-GBM) glioma (A) and GBM (B) tissue. Original amplification was 4x.

